# Additive effects of cerebrovascular disease functional connectome phenotype and plasma p-tau181 on longitudinal neurodegeneration and cognitive outcomes

**DOI:** 10.1101/2024.07.09.602637

**Authors:** Joanna Su Xian Chong, Fang Ji, Saima Hilal, Joyce Ruifen Chong, Jia Ming Lau, Nathanael Ren Jie Tong, Boon Yeow Tan, Narayanaswamy Venketasubramanian, Mitchell Kim Peng Lai, Christopher Li-Hsian Chen, Juan Helen Zhou

## Abstract

**INTRODUCTION:** We investigated the effects of multiple cerebrovascular disease (CeVD) neuroimaging markers on brain functional connectivity (FC), and how such CeVD-related FC changes interact with plasma p-tau181 (Alzheimer’s disease (AD) marker) to influence downstream neurodegeneration and cognitive changes.

**METHODS:** Multivariate associations between four CeVD markers and whole-brain FC in 529 participants across the dementia spectrum were examined using partial least squares correlation. Interactive effects of CeVD-related FC patterns and p-tau181 on longitudinal grey matter volume and cognitive changes were investigated using linear mixed-effects models.

**RESULTS:** We identified a brain FC phenotype associated with high CeVD burden across all markers. Further, expression of this general CeVD-related FC phenotype and p-tau181 contributed additively, but not synergistically, to baseline and longitudinal grey matter volumes and cognitive changes.

**DISCUSSION:** Our findings suggest that CeVD exerts global effects on the brain connectome and highlight the additive nature of AD and CeVD on neurodegeneration and cognition.

## 1 INTRODUCTION

Cerebrovascular disease (CeVD) is a major cause of cognitive impairment in older adults, being second only to Alzheimer’s disease (AD) as the most common contributor to dementia [1]. CeVD is identified by several magnetic resonance imaging (MRI) markers such as white matter hyperintensities, lacunes and microbleeds [2], and is associated with multiple clinical symptoms, including cognitive decline across multiple domains [3], neuropsychiatric symptoms [4,5], mood disturbances [6] and movement impairments [7]. Further, AD and CeVD are frequently co-morbid in individuals with cognitive impairment, with up to 75% of AD cases showing co-occurring CeVD pathology at autopsy [8]. Consequently, understanding how CeVD affects the brain and interacts with AD to contribute to cognitive decline is an important research goal.

In recent years, functional MRI (fMRI)-derived measures of intrinsic functional connectivity (FC), which measures the temporal synchrony between low-frequency fluctuations of blood-oxygenation-level-dependent (BOLD) signals in brain regions under resting-state conditions, have provided valuable insight on how CeVD gives rise to diverse clinical symptoms beyond its focal lesions through its effect on the functional connectome [9,10]. Studies have reported disrupted FC across large-scale brain networks in individuals with CeVD, particularly in the executive control, dorsal attention, salience and default mode networks [9,10], which are in turn linked to poorer cognitive function [11,12]. These widespread CeVD-related effects on FC thus reinforce the idea of CeVD being a global rather than a focal disease [10]. Additionally, converging evidence from pathological, neuroimaging and clinical studies have suggested that AD and CeVD have additive, rather than synergistic, effects on cognitive function [13–17]. Autopsy studies, for example, have reported lower AD burden in AD patients with CeVD compared to those without CeVD given the same dementia severity [14]. Correspondingly, our previous work has demonstrated divergent effects of AD and CeVD burden on FC [15,16]. In a study on amnestic and subcortical vascular mild cognitive impairment patients, we showed that amyloid-β burden was associated with longitudinal default mode network FC disruptions, while baseline lacune number was linked to longitudinal executive control network FC disruptions [15].

One major limitation of these fMRI studies is that CeVD burden is typically quantified using a single MRI marker (e.g., lacune numbers) when examined in a continuous manner [9]. However, CeVD encompasses several heterogeneous MRI markers that vary in aetiology and disease severity [2,10]. As such, the extent of CeVD burden may not be fully reflected using a single marker. A multivariate approach that accounts for multiple CeVD markers could hence better reflect the influence of CeVD on FC and its subsequent effects on cognitive decline. Such an approach would also enable investigations into whether different CeVD markers are linked to differential changes in FC, or whether CeVD markers collectively exert a global effect on FC.

Our study thus sought to examine the effects of various CeVD MRI markers on whole-brain FC using a multivariate approach in individuals at different cognitive stages and with varying degrees of AD and CeVD burden. Given that CeVD has been suggested to be a global rather than a focal disease due to its widespread effects on the brain connectome [10], we hypothesized that CeVD would be linked to widespread FC changes in a non-marker specific manner. Further, we examined how such FC disruptions related to multiple CeVD markers would interact with AD pathology to influence baseline and longitudinal changes in downstream outcomes such as cognitive/behavioural performance and grey matter volumes. As a marker of AD pathology, we used plasma p-tau181, an emerging biomarker with growing evidence demonstrating its strong association with positron emission tomography amyloid-β uptake, as well as its ability to predict AD progression and distinguish AD from other neurodegenerative diseases [18–20]. Given evidence suggesting additive effects of CeVD and AD on cognition [13–17], we hypothesized that expression of the CeVD-related FC phenotype and p-tau181 would have additive rather than synergistic effects on cognitive/behavioural performance and grey matter volume changes at baseline and over time.

## 2 METHODS

### 2.1 Participants

700 participants with no cognitive impairment (NCI), cognitive impairment no dementia (CIND) or dementia were recruited from memory clinics at National University Hospital, Singapore and St. Luke’s Hospital, Singapore (NCI, CIND and dementia participants), as well as from the community (NCI participants only). At baseline, all participants underwent a comprehensive protocol comprising physical, clinical, neuroimaging and neuropsychological assessments, including a locally validated formal neuropsychological test battery. Cognitive assessments were performed annually and neuroimaging assessments were repeated after 2 and 4 to 5 years. The study was approved by the Singapore National Healthcare Group Domain-Specific Review Board and written informed consent was obtained from all participants in their preferred language before study commencement.

NCI was defined as having no cognitive impairment in all domains on the neuropsychological test battery. CIND was defined as having an impairment in at least one cognitive domain on the neuropsychological test battery, but not meeting the diagnosis for dementia. Dementia was defined based on the Diagnostic and Statistical Manual of Mental Disorders, Fourth Edition (DSM-IV), with aetiological diagnoses of AD made using the National Institute of Neurological and Communicative Disorders and Stroke and the Alzheimer’s Disease and Related Disorders Association (NINCDS-ADRDA) [21] and diagnoses of vascular dementia/mixed dementia made using the National Institute of Neurological Disorders and Stroke and Association Internationale pour la Recherche et l’ Enseignement en Neurosciences (NINDS-AIREN) criteria [22].

For this study, we considered only participants with good quality multimodal neuroimaging data and complete set of CeVD visual ratings at baseline. In total, 529 participants with at least four CeVD features (number of lacunes, number of cortical infarcts, age-related white matter changes (ARWMC) and number of cerebral microbleeds) were included in the study (see Table 1 for participants’ demographic information and Supplementary Figure 1 for the participant flowchart).

**Table 1:**
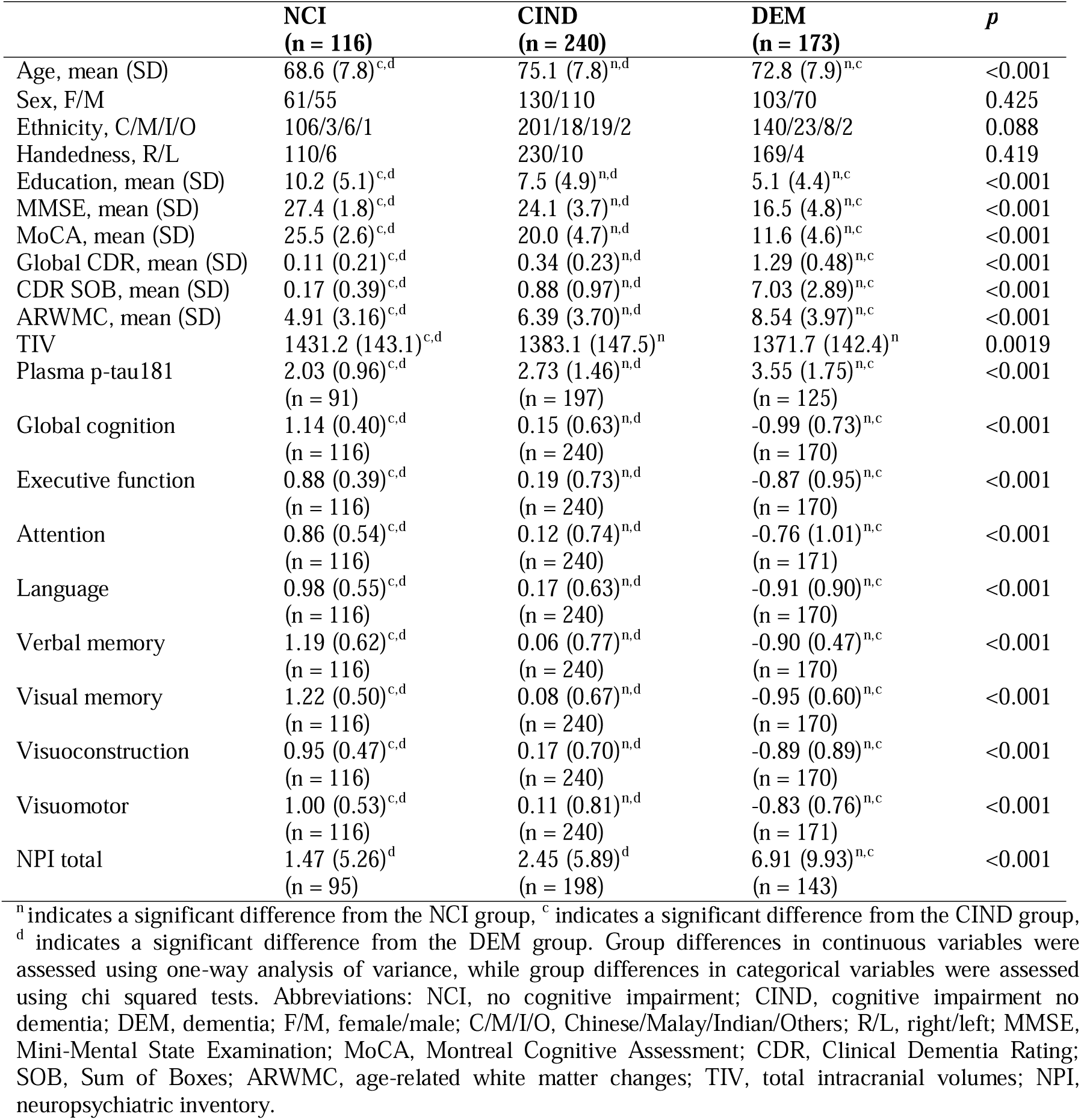
Participant demographic and clinical characteristics at baseline.

|  | **NCI**<br>**(n = 116)** | **CIND**<br>**(n = 240)** | **DEM**<br>**(n = 173)** | **p** |
| --- | --- | --- | --- | --- |
| Age, mean (SD) | 68.6 (7.8)<sup>c,d</sup> | 75.1 (7.8)<sup>n,d</sup> | 72.8 (7.9)<sup>n,c</sup> | <0.001 |
| Sex, F/M | 61/55 | 130/110 | 103/70 | 0.425 |
| Ethnicity, C/M/I/O | 106/3/6/1 | 201/18/19/2 | 140/23/8/2 | 0.088 |
| Handedness, R/L | 110/6 | 230/10 | 169/4 | 0.419 |
| Education, mean (SD) | 10.2 (5.1)<sup>c,d</sup> | 7.5 (4.9)<sup>n,d</sup> | 5.1 (4.4)<sup>n,c</sup> | <0.001 |
| MMSE, mean (SD) | 27.4 (1.8)<sup>c,d</sup> | 24.1 (3.7)<sup>n,d</sup> | 16.5 (4.8)<sup>n,c</sup> | <0.001 |
| MoCA, mean (SD) | 25.5 (2.6)<sup>c,d</sup> | 20.0 (4.7)<sup>n,d</sup> | 11.6 (4.6)<sup>n,c</sup> | <0.001 |
| Global CDR, mean (SD) | 0.11 (0.21)<sup>c,d</sup> | 0.34 (0.23)<sup>n,d</sup> | 1.29 (0.48)<sup>n,c</sup> | <0.001 |
| CDR SOB, mean (SD) | 0.17 (0.39)<sup>c,d</sup> | 0.88 (0.97)<sup>n,d</sup> | 7.03 (2.89)<sup>n,c</sup> | <0.001 |
| ARWMC, mean (SD) | 4.91 (3.16)<sup>c,d</sup> | 6.39 (3.70)<sup>n,d</sup> | 8.54 (3.97)<sup>n,c</sup> | <0.001 |
| TIV | 1431.2 (143.1)<sup>c,d</sup> | 1383.1 (147.5)<sup>n</sup> | 1371.7 (142.4)<sup>n</sup> | 0.0019 |
| Plasma p-tau181 | 2.03 (0.96)<sup>c,d</sup> | 2.73 (1.46)<sup>n,d</sup> | 3.55 (1.75)<sup>n,c</sup> | <0.001 |
|  | (n = 91) | (n = 197) | (n = 125) |  |
| Global cognition | 1.14 (0.40)<sup>c,d</sup> | 0.15 (0.63)<sup>n,d</sup> | -0.99 (0.73)<sup>n,c</sup> | <0.001 |
|  | (n = 116) | (n = 240) | (n = 170) |  |
| Executive function | 0.88 (0.39)<sup>c,d</sup> | 0.19 (0.73)<sup>n,d</sup> | -0.87 (0.95)<sup>n,c</sup> | <0.001 |
|  | (n = 116) | (n = 240) | (n = 170) |  |
| Attention | 0.86 (0.54)<sup>c,d</sup> | 0.12 (0.74)<sup>n,d</sup> | -0.76 (1.01)<sup>n,c</sup> | <0.001 |
|  | (n = 116) | (n = 240) | (n = 171) |  |
| Language | 0.98 (0.55)<sup>c,d</sup> | 0.17 (0.63)<sup>n,d</sup> | -0.91 (0.90)<sup>n,c</sup> | <0.001 |
|  | (n = 116) | (n = 240) | (n = 170) |  |
| Verbal memory | 1.19 (0.62)<sup>c,d</sup> | 0.06 (0.77)<sup>n,d</sup> | -0.90 (0.47)<sup>n,c</sup> | <0.001 |
|  | (n = 116) | (n = 240) | (n = 170) |  |
| Visual memory | 1.22 (0.50)<sup>c,d</sup> | 0.08 (0.67)<sup>n,d</sup> | -0.95 (0.60)<sup>n,c</sup> | <0.001 |
|  | (n = 116) | (n = 240) | (n = 170) |  |
| Visuoconstruction | 0.95 (0.47)<sup>c,d</sup> | 0.17 (0.70)<sup>n,d</sup> | -0.89 (0.89)<sup>n,c</sup> | <0.001 |
|  | (n = 116) | (n = 240) | (n = 170) |  |
| Visuomotor | 1.00 (0.53)<sup>c,d</sup> | 0.11 (0.81)<sup>n,d</sup> | -0.83 (0.76)<sup>n,c</sup> | <0.001 |
|  | (n = 116) | (n = 240) | (n = 171) |  |
| NPI total | 1.47 (5.26)<sup>d</sup> | 2.45 (5.89)<sup>d</sup> | 6.91 (9.93)<sup>n,c</sup> | <0.001 |
|  | (n = 95) | (n = 198) | (n = 143) |  |

<sup>n</sup> indicates a significant difference from the NCI group, <sup>c</sup> indicates a significant difference from the CIND group, <sup>d</sup> indicates a significant difference from the DEM group. Group differences in continuous variables were assessed using one-way analysis of variance, while group differences in categorical variables were assessed using chi squared tests. Abbreviations: NCI, no cognitive impairment; CIND, cognitive impairment no dementia; DEM, dementia; F/M, female/male; C/M/I/O, Chinese/Malay/Indian/Others; R/L, right/left; MMSE, Mini-Mental State Examination; MoCA, Montreal Cognitive Assessment; CDR, Clinical Dementia Rating; SOB, Sum of Boxes; ARWMC, age-related white matter changes; TIV, total intracranial volumes; NPI, neuropsychiatric inventory.

### 2.2 Neuropsychological assessment

Participants completed a comprehensive neuropsychological assessment administered by trained research psychologists, which included the Mini-Mental State Examination (MMSE) [23], Montreal Cognitive Assessment (MoCA) [24], Clinical Dementia Rating (CDR) [25], Neuropsychiatric Inventory (NPI) [26], as well as a formal neuropsychological test battery locally validated in elderly Singaporeans [27]. For the NPI, a total score was obtained by multiplying the frequency and severity scores for each symptom (resulting in a score for each symptom), then summing all individual symptom scores together.

The neuropsychological test battery assessed seven cognitive domains (five non-memory, two memory domains) using the following tests: (1) executive function: frontal assessment battery [28] and maze task [29]; (2) attention: digit span, visual memory span [30] and auditory detection tests [31], (3) language: Boston naming test [32] and verbal fluency [33]; (4) visuomotor speed: symbol digit modalities test [34] and digit cancellation [35]; (5) visuoconstruction: Wechsler Memory Scale-Revised (WMS-R) visual reproduction copy task [30], clock drawing [36] and Wechsler Adult Intelligence Scale-Revised (WAIS-R) subtest of block design [37]; (6) verbal memory: word list recall [38] and story recall; and (7) visual memory: picture recall and WMS-R visual reproduction task [30]. For each domain, a composite z-score for each participant was calculated as follows: 1) transformation of each test’s raw scores to z-scores using the mean and standard deviation of scores in the current sample; 2) averaging of z-scores of all tests in each domain to obtain domain-specific z-scores; and 3) standardization of domain-specific z-scores using the mean and standard deviation of domain-specific z-scores in the current sample. Additionally, we obtained a measure of global cognition by averaging across all domain-specific z-scores and subsequently standardizing the global z-scores using the mean and standard deviation of global z-scores in the sample.

### 2.3 Plasma p-tau181 measurements

Non-fasting blood was collected in ethylenediaminetetraacetic acid (EDTA) tubes and centrifuged at 2000 rcf for 10 minutes at 4°C. Following centrifugation, plasma was extracted, mixed well, aliquoted into polypropylene tubes (0.2 ml per tube) and stored at −80°C until use. Plasma p-tau181 was measured on the Simoa HD-1 Analyzer using the commercially available Simoa pTau-181 Advantage V2 kit per manufacturer’s instructions (Quanterix, Billerica, Massachusetts, USA).

### 2.4 Image acquisition

Participants underwent neuroimaging scans at the Clinical Imaging Research Centre, National University of Singapore, on a 3T Siemens Magnetom Tim Trio scanner using a 32-channel head coil. The scan protocol included T1-weighted, T2-weighted, fluid attenuated inversion fecovery (FLAIR), susceptibility weighted imaging (SWI), magnetic resonance angiography (MRA) and T2*-weighted resting state fMRI sequences.

T1-weighted scans were acquired using a magnetization prepared rapid gradient echo (MPRAGE) sequence (repetition time = 2300 ms, echo time = 1.9 ms, inversion time = 900 ms, flip angle = 9°, voxel size = 1.0 mm isotropic, field of view = 256 × 256 mm^2^, 192 sagittal slices), while T2*-weighted resting state fMRI scans were acquired using an echo planar sequence (acquisition time = 5 min, repetition time = 2300 ms, echo time = 25 ms, flip angle = 90°, voxel size = 3.0 mm isotropic, field of view = 192 × 192 mm^2^, 48 sagittal slices, interleaved collection). Protocols for the full set of scans performed during the study, including those used for assessment of CeVD markers, are described in previous work [39].

### 2.5 Image preprocessing

#### 2.5.1 Functional imaging

Resting state fMRI images were preprocessed following previous procedures [16] using the FMRIB (Oxford Centre for Functional MRI of the Brain) Software Library (FSL) [40] and Analysis of Functional NeuroImages (AFNI) software [41]. The preprocessing steps comprised: 1) discarding the first five volumes to stabilize the magnetic field; 2) motion and slice timing correction; 3) despiking of time series; 4) grand mean scaling; 5) spatial smoothing using a 6 mm Gaussian kernel, 6) temporal band-pass filtering (0.009 – 0.1 Hz); 7) linear and quadratic detrending; 8) co-registration of fMRI image to T1-weighted image using boundary based registration followed by non-linear registration (FNIRT) of fMRI image to standard Montreal Neurological Institute 152 (MNI152) space; and 9) regression of nuisance signals (white matter, cerebrospinal fluid, global signal and six motion parameters).

#### 2.5.2 Structural imaging

Voxel-based morphometry (VBM) was performed on T1-weighted images to obtain voxel-wise grey matter volume probability maps, using the computational anatomy toolbox (CAT12 v12.7) [42] for Statistical Parametric Mapping (SPM12; Wellcome Trust Centre for Neuroimaging; http://www.fil.ion.ucl.ac.uk/spm/software/spm12/). The longitudinal VBM pipeline optimized for capturing larger changes over time (such as atrophy due to AD or ageing) was used, and included the following steps: 1) intra-participant (across time points) bias field correction and affine registration of T1-weighted images to create participant-specific mean images across time points; 2) segmentation of the resultant mean images into grey matter, white matter and cerebrospinal fluid using the standard CAT12 processing pipeline to obtain participant-specific tissue probability maps, which were subsequently used to enhance the time point-specific registrations to create the final grey matter, white matter and cerebrospinal fluid segments; 4) estimation of deformation parameters of tissue segments to a study-specific template in MNI space (created using baseline T1-weighted images of all participants in the cohort) using DARTEL (Diffeomorphic Anatomical Registration Through Exponentiated Lie Algebra) registration while also taking into account deformations between images of different time points to account for age-related changes over time; 5) application of mean deformation across all time points to individual grey and white matter segmented images; and 6) modulation of voxel values with both the linear and non-linear components of the Jacobian determinant to enable comparison of absolute tissue volumes.

#### 2.5.3 Assessment of cerebrovascular disease markers

MRI markers of CeVD were graded using the Standards for Reporting Vascular changes on Neuroimaging (STRIVE) criteria [2]. Lacunes were defined as subcortical lesions, 3 to 15 mm in diameter, with high signal on T2-weighted images and low signal on T1-weighted and FLAIR images, and a hyperintense rim with a centre following the cerebrospinal fluid intensity [2]. Cerebral microbleeds were defined as focal, round hypointense lesions with blooming effect on SWI images and graded using the Brain Observer Microbleed Scale [43]. White matter hyperintensities on T2-weighted and FLAIR images were graded using the ARWMC scale [44]. Cortical infarcts were defined as focal lesions involving cortical grey matter, with size > 5 mm in diameter, hyperintense rim on FLAIR images and centre following cerebrospinal fluid intensity, as well as tissue loss of variable magnitude with prominent adjacent sulci and ipsilateral ventricular enlargement [2]. Finally, cortical cerebral microinfarcts were defined as cortical lesions < 5 mm in diameter and perpendicular to the cortical surface, with hypointense signal on T1-weighted images and hyperintense signal on T2-weighted and FLAIR images [45].

#### 2.5.4 Derivation of functional connectivity matrices and regional grey matter volumes

Undirected, weighted FC matrices were derived for each participant. Mean BOLD time series of 144 regions-of-interest (ROIs) interest (114 cortical regions [46], 30 subcortical regions [47,48]) were first extracted from preprocessed fMRI images in MNI152 space. Pearson correlations between the mean BOLD time series of each and every pair of ROIs were then computed and transformed to z-scores using Fisher’s r-to-z transformation, resulting in a 144 × 144 FC matrix. For the analyses, we computed network-level 20 × 20 FC matrices describing the connectivity within– and between-20 subnetworks (see Supplementary Table 1 for the list of networks) by further averaging across connections of all ROIs within each network as well as between pairs of networks.

Regional grey matter volumes for each participant were obtained by averaging grey matter volumes of all voxels within each of the 144 ROIs. For statistical analyses, we further derived grey matter volumes of 10 networks (default, control, limbic, salience/ventral attention, dorsal attention, somatomotor, visual, temporoparietal, hippocampal and subcortical) by averaging across grey matter volumes of all ROIs belonging to each network.

### 2.6 Statistical analyses

#### 2.6.1 Multivariate associations between cerebrovascular disease markers and functional connectivity

Partial least squares correlation (PLSC) was used to examine multivariate associations between the network-level 20 × 20 (i.e., (20 × 19 / 2) = 190 connections) FC matrices and 4 CeVD markers (number of lacunes, number of cerebral microbleeds, number of cortical infarcts and ARWMC). PLSC is a multivariate approach which uses singular vector decomposition to identify a set of latent variables (i.e., linear combinations of the original variables) that maximises the covariance between two sets of variables [49]. Compared to multiple regression, PLSC is able to more robustly handle highly correlated observations, as well as datasets where the number of predictors exceed the number of samples (i.e., p >> n problem) [50]. A schematic of the multivariate analyses is provided in Figure 1. First, nuisance covariates (age, sex, ethnicity, years of education, handedness and total intracranial volumes (TIV)) were regressed out from the FC matrices and log-transformed CeVD markers (log transformation was done due to the skewness of the values) using multiple regression models. Behaviour PLSC was then performed on the standardized FC and CeVD residuals using the PLS toolbox [51] in MATLAB. The statistical significance of the latent variables was determined using permutation tests (1000 permutations, significance reached at *p* < 0.05), while the reliability of each functional connection’s contribution (i.e., connectome weight) to the latent variable was established using bootstrap ratios, which is computed by dividing the weight by the standard error estimated from 100 bootstrap samples. Bootstrap ratios are analogous to z-scores, and functional connections with ratios > 2 are considered to be significantly stable and reliable [49]. For CeVD markers, the reliability of their weights in the latent variable was determined using the bootstrapped confidence intervals of the correlation between connectome scores and each of the CeVD variables [49,51].

**Figure 1:**
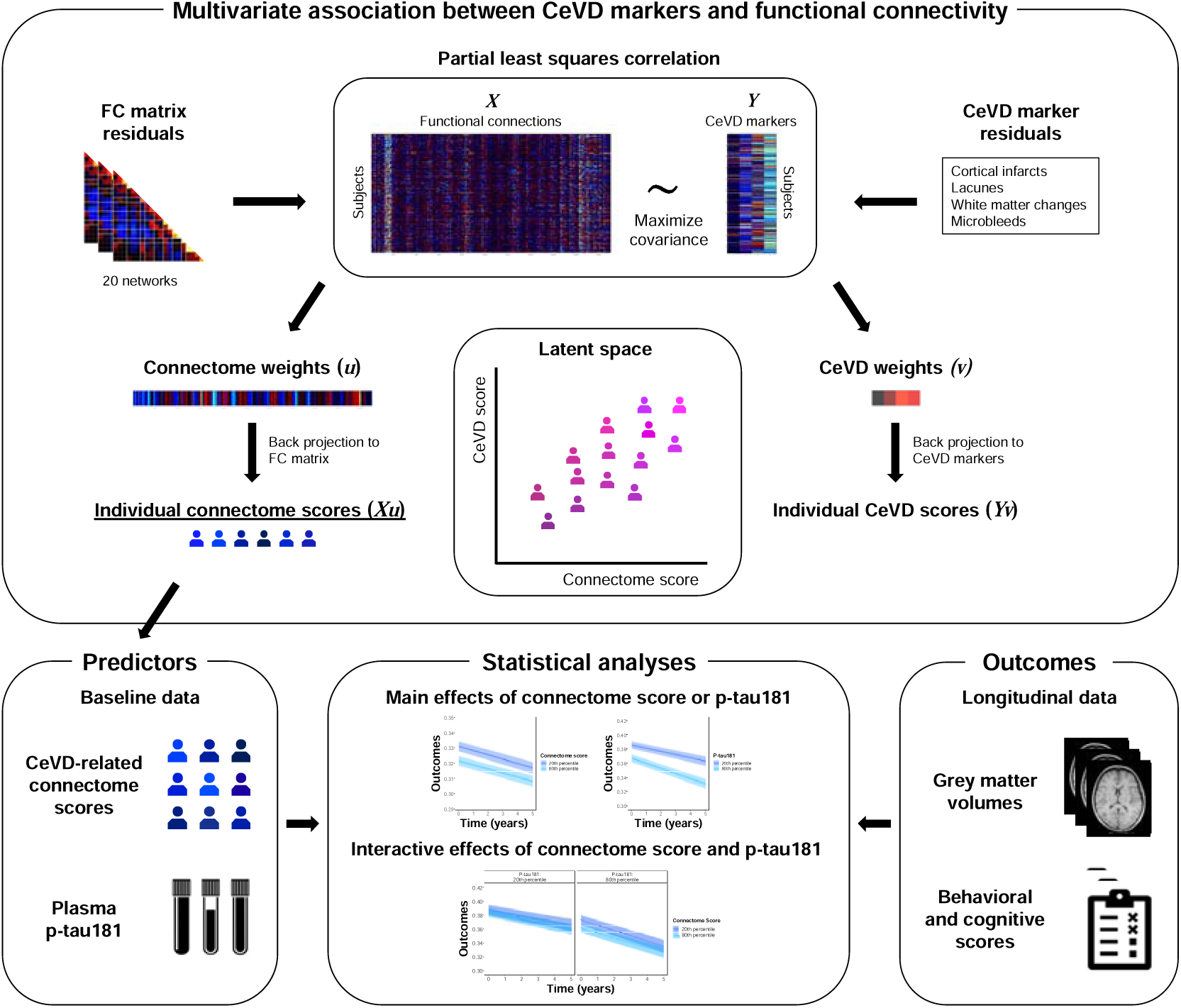
Study design schematic. Multivariate associations between residuals (after regressing out nuisance variables) of whole-brain FC and multiple CeVD markers were examined using partial least squares correlation, an approach which seeks to identify a set of latent variables (i.e., linear combinations of the original variables) that maximizes the covariance between two sets of variables. The resultant FC *(u)* and CeVD *(v)* weights were then respectively back projected to their original residual values to obtain connectome *(Xu)* and CeVD *(Yv)* scores for each participant. Connectome scores, in particular, denote the extent to which each participant expressed the FC pattern that is maximally related to the CeVD markers. Main and interactive effects of connectome score and plasma p-tau181 (an AD marker) on downstream outcomes, including grey matter volumes and cognitive/behavioural performance, were subsequently analyzed using linear mixed effects models. Abbreviations: FC, functional connectivity; CeVD, cerebrovascular disease; AD, Alzheimer’s disease.

#### 2.6.2 Validation of the cerebrovascular disease-related functional connectome phenotype

To examine the reproducibility of the CeVD-linked FC pattern in the original analyses, we repeated the PLSC analyses on two separate datasets: 1) baseline data of a subset of participants with five CeVD features (number of lacunes, number of cortical infarcts, ARWMC, number of cerebral microbleeds and number of cortical microinfarcts) (n = 436 comprising 93 NCI, 198 CIND and 145 dementia), and 2) year 2 data (i.e., 2 years from baseline) of participants with four CeVD features (n = 316 comprising 96 NCI, 119 CIND, 101 dementia) (see Supplementary Tables 2 and 3 for demographic characteristics of these datasets).

#### 2.6.3 Main and interactive effects of cerebrovascular disease-related connectome scores and p-tau181 on downstream variables

We next sought to examine how this CeVD-related FC pattern was linked to AD pathology (p-tau181), and whether they interact to influence downstream variables such as behavioural/cognitive performance and grey matter volumes. To this end, CeVD-related connectome scores were obtained from the PLSC analyses by back projecting the connectome weights to the original FC residuals (i.e., computing the dot product of FC residuals and connectome weights). Connectome scores describe the extent to which each participant expressed this CeVD-related FC phenotype, with higher connectome scores indicating greater expression of the FC pattern. We first examined the association between log-transformed p-tau181 (log transformation was done due to the skewness of the p-tau181 values) and connectome scores using linear regression (n = 413), controlling for age, sex, ethnicity, years of education, handedness, TIV and diagnosis. Next, we examined main and interactive effects of baseline p-tau181 and connectome scores on baseline and longitudinal changes in behavioural/cognitive performance and grey matter volumes. To accomplish this, linear mixed effects models were run in a subset of participants with baseline plasma p-tau181 and longitudinal neuropsychological and neuroimaging data (see Tables 2 and 3 for sample sizes). In these models, time was modelled as a random effect, and age, sex, ethnicity, years of education, handedness, TIV, diagnosis, p-tau181, and connectome score were modelled as fixed effects, as follows:

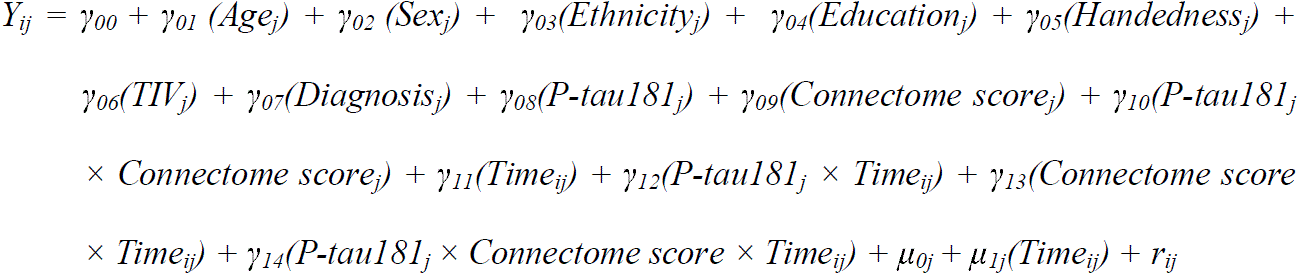

**Table 2:** Main and interaction effects of connectome score and p-tau181 on baseline and longitudinal changes in grey matter volumes.

| **Measure** | **Predictor** | **n** | **Coeff.** | **SE** | **t** | **Adj. p** |
| --- | --- | --- | --- | --- | --- | --- |
| Default | Connectome score | NCI: 80 | -1.04e<sup>-2</sup> | 2.87e<sup>-3</sup> | -3.62 | 0.0017* |
|  | P-tau181 | CIND: 155 | -1.00e<sup>-2</sup> | 4.13e<sup>-3</sup> | -2.42 | 0.0402* |
|  | Connectome score × P-tau181 | DEM: 81 | 5.13e<sup>-3</sup> | 8.27e<sup>-3</sup> | 0.62 | 0.9936 |
|  | Connectome score × Time |  | 5.38e<sup>-5</sup> | 3.48e<sup>-4</sup> | 0.15 | 0.9785 |
|  | P-tau181 × Time |  | -1.67e<sup>-3</sup> | 4.57e<sup>-4</sup> | -3.66 | 0.0008* |
|  | Connectome score × P-tau181 |  | -3.94e<sup>-4</sup> | 1.04e<sup>-3</sup> | -0.38 | 0.7190 |

|  | × Time |  |  |  |  |  |
| --- | --- | --- | --- | --- | --- | --- |
| Control | Connectome score | NCI: 80 | -1.32e<sup>-2</sup> | 2.80e<sup>-3</sup> | -4.70 | <0.0001* |
|  | P-tau181 | CIND: 155 | -9.39e<sup>-3</sup> | 4.03e<sup>-3</sup> | -2.33 | 0.0403* |
|  | Connectome score × P-tau181 | DEM: 81 | 5.49e<sup>-3</sup> | 8.06e<sup>-3</sup> | 0.68 | 0.9936 |
|  | Connectome score × Time |  | 1.70e<sup>-4</sup> | 3.36e<sup>-4</sup> | 0.51 | 0.9785 |
|  | P-tau181 × Time |  | -1.34e<sup>-3</sup> | 4.42e<sup>-4</sup> | -3.03 | 0.0051* |
|  | Connectome score × P-tau181 × Time |  | -3.89e<sup>-4</sup> | 1.01e<sup>-3</sup> | -0.39 | 0.7190 |
| Limbic | Connectome score | NCI: 80 | -1.14e<sup>-2</sup> | 3.90e<sup>-3</sup> | -2.93 | 0.0061* |
|  | P-tau181 | CIND: 155 | -1.27e<sup>-2</sup> | 5.61e<sup>-3</sup> | -2.27 | 0.0403* |
|  | Connectome score × P-tau181 | DEM: 81 | 7.95e<sup>-5</sup> | 1.12e<sup>-2</sup> | 0.01 | 0.9944 |
|  | Connectome score × Time |  | 3.73e<sup>-4</sup> | 4.69e<sup>-4</sup> | 0.80 | 0.9530 |
|  | P-tau181 × Time |  | -3.30e<sup>-3</sup> | 6.18e<sup>-4</sup> | -5.34 | <0.0001* |
|  | Connectome score × P-tau181 × Time |  | 9.66e<sup>-4</sup> | 1.39e<sup>-3</sup> | 0.69 | 0.7190 |
| Salience/ventral attention | Connectome score | NCI: 80 | -6.97e<sup>-3</sup> | 2.84e<sup>-3</sup> | -2.45 | 0.0185* |
|  | P-tau181 | CIND: 155 | -1.84e<sup>-3</sup> | 4.09e<sup>-3</sup> | -0.45 | 0.6528 |
|  | Connectome score × P-tau181 | DEM: 81 | 1.50e<sup>-3</sup> | 8.19e<sup>-3</sup> | 0.18 | 0.9944 |
|  | Connectome score × Time |  | 2.62e<sup>-4</sup> | 3.67e<sup>-4</sup> | 0.71 | 0.9530 |
|  | P-tau181 × Time |  | -1.44e<sup>-3</sup> | 4.81e<sup>-4</sup> | -2.99 | 0.0051* |
|  | Connectome score × P-tau181 × Time |  | -5.63e<sup>-4</sup> | 1.10e<sup>-3</sup> | -0.51 | 0.7190 |
| Dorsal attention | Connectome score | NCI: 80 | -8.68e<sup>-3</sup> | 2.82e<sup>-3</sup> | -3.08 | 0.0046* |
|  | P-tau181 | CIND: 155 | -7.63e<sup>-3</sup> | 4.06e<sup>-3</sup> | -1.88 | 0.0872 |
|  | Connectome score × P-tau181 | DEM: 81 | 2.02e<sup>-3</sup> | 8.12e<sup>-3</sup> | 0.25 | 0.9944 |
|  | Connectome score × Time |  | 1.00e<sup>-4</sup> | 3.65e<sup>-4</sup> | 0.28 | 0.9785 |
|  | P-tau181 × Time |  | -1.82e<sup>-3</sup> | 4.79e<sup>-4</sup> | -3.79 | 0.0006* |
|  | Connectome score × P-tau181 × Time |  | -6.66e<sup>-4</sup> | 1.09e<sup>-3</sup> | -0.61 | 0.7190 |
| Somatomotor | Connectome score | NCI: 80 | -9.16e<sup>-3</sup> | 2.93e<sup>-3</sup> | -3.13 | 0.0046* |
|  | P-tau181 | CIND: 155 | 2.65e<sup>-3</sup> | 4.21e<sup>-3</sup> | 0.63 | 0.6528 |
|  | Connectome score × P-tau181 | DEM: 81 | 4.79e<sup>-4</sup> | 8.43e<sup>-3</sup> | 0.06 | 0.9944 |
|  | Connectome score × Time |  | 3.42e<sup>-4</sup> | 3.55e<sup>-4</sup> | 0.96 | 0.9530 |
|  | P-tau181 × Time |  | -6.84e<sup>-4</sup> | 4.64e<sup>-4</sup> | -1.47 | 0.1684 |
|  | Connectome score × P-tau181 × Time |  | -8.70e<sup>-4</sup> | 1.07e<sup>-3</sup> | -0.81 | 0.7190 |
| Visual | Connectome score | NCI: 80 | -9.20e<sup>-3</sup> | 2.98e<sup>-3</sup> | -3.09 | 0.0046* |
|  | P-tau181 | CIND: 155 | -1.04e<sup>-2</sup> | 4.28e<sup>-3</sup> | -2.42 | 0.0402* |
|  | Connectome score × P-tau181 | DEM: 81 | 7.51e<sup>-3</sup> | 8.58e<sup>-3</sup> | 0.87 | 0.9936 |
|  | Connectome score × Time |  | 2.65e<sup>-4</sup> | 3.48e<sup>-4</sup> | 0.76 | 0.9530 |
|  | P-tau181 × Time |  | -1.03e<sup>-3</sup> | 4.54e<sup>-4</sup> | -2.27 | 0.0348* |
|  | Connectome score × P-tau181 × Time |  | -3.79e<sup>-4</sup> | 1.05e<sup>-3</sup> | -0.36 | 0.7190 |
| Temporoparietal | Connectome score | NCI: 80 | -8.29e<sup>-3</sup> | 4.46e<sup>-3</sup> | -1.86 | 0.0713 |
|  | P-tau181 | CIND: 155 | -2.85e<sup>-2</sup> | 6.42e<sup>-3</sup> | -4.44 | 0.0001* |
|  | Connectome score × P-tau181 | DEM: 81 | 6.81e<sup>-3</sup> | 1.28e<sup>-2</sup> | 0.53 | 0.9936 |
|  | Connectome score × Time |  | -1.23e<sup>-5</sup> | 4.55e<sup>-4</sup> | -0.03 | 0.9785 |
|  | P-tau181 × Time |  | -8.53e<sup>-4</sup> | 5.93e<sup>-4</sup> | -1.44 | 0.1684 |
|  | Connectome score × P-tau181 × Time |  | 1.22e<sup>-3</sup> | 1.37e<sup>-3</sup> | 0.89 | 0.7190 |
| Hippocampus | Connectome score | NCI: 80 | -1.06e<sup>-2</sup> | 4.27e<sup>-3</sup> | -2.49 | 0.0185* |
|  | P-tau181 | CIND: 155 | -3.01e<sup>-2</sup> | 6.12e<sup>-3</sup> | -4.91 | <0.0001* |
|  | Connectome score × P-tau181 | DEM: 81 | -1.85e<sup>-2</sup> | 1.23e<sup>-2</sup> | -1.50 | 0.9936 |
|  | Connectome score × Time |  | 2.86e<sup>-5</sup> | 5.13e<sup>-4</sup> | 0.06 | 0.9785 |
|  | P-tau181 × Time |  | -4.15e<sup>-3</sup> | 6.76e<sup>-4</sup> | -6.14 | <0.0001* |
|  | Connectome score × P-tau181 × Time |  | 2.42e<sup>-3</sup> | 1.53e<sup>-3</sup> | 1.58 | 0.7190 |
|  | × Time |  |  |  |  |  |
| Subcortical | Connectome score | NCI: 80 | -4.87e<sup>-3</sup> | 5.27e<sup>-3</sup> | -0.92 | 0.3564 |
|  | P-tau181 | CIND: 155 | -3.78e<sup>-3</sup> | 7.57e<sup>-3</sup> | -0.50 | 0.6528 |
|  | Connectome score × P-tau181 | DEM: 81 | -1.50e<sup>-2</sup> | 1.52e<sup>-2</sup> | -0.99 | 0.9936 |
|  | Connectome score × Time |  | -1.07e<sup>-3</sup> | 8.00e<sup>-4</sup> | -1.33 | 0.9530 |
|  | P-tau181 × Time |  | -1.23e<sup>-3</sup> | 1.05e<sup>-3</sup> | -1.17 | 0.2438 |
|  | Connectome score × P-tau181 |  | 3.31e<sup>-3</sup> | 2.41e<sup>-3</sup> | 1.38 | 0.7190 |
|  | × Time |  |  |  |  |  |

\* indicates significance at $p < 0.05$ , after false discovery rate corrections for multiple comparisons across the ten brain networks. Abbreviations: Coeff., coefficient; SE, standard error; Adj. $p$ , $p$ -value adjusted for false discovery rate; NCI, no cognitive impairment; CIND, cognitive impairment no dementia; DEM, dementia.

**Table 3:** Main and interaction effects of connectome score and p-tau181 on baseline and longitudinal changes in cognitive and behavioural performance.

| Measure | Predictor | n | Coeff. | SE | t | Adj. $p$ |
| --- | --- | --- | --- | --- | --- | --- |
| CDR sum of boxes | Connectome score | NCI: 77 | 0.62 | 0.20 | 3.18 | 0.0050* |
|  | P-tau181 | CIND: 148 | 0.55 | 0.28 | 1.93 | 0.0815 |
|  | Connectome score × P-tau181 | DEM: 83 | 0.76 | 0.56 | 1.37 | 0.9255 |
|  | Connectome score × Time |  | 0.39 | 0.11 | 3.40 | 0.0094* |
|  | P-tau181 × Time |  | 1.01 | 0.15 | 6.66 | <0.0001* |
|  | Connectome score × P-tau181 |  | -0.33 | 0.33 | -0.99 | 0.6953 |
|  | × Time |  |  |  |  |  |
| MMSE | Connectome score | NCI: 76 | -1.41 | 0.40 | -3.50 | 0.0022* |
|  | P-tau181 | CIND: 147 | -2.13 | 0.58 | -3.67 | 0.0009* |
|  | Connectome score × P-tau181 | DEM: 85 | 0.05 | 1.12 | 0.05 | 0.9631 |
|  | Connectome score × Time |  | -0.18 | 0.13 | -1.38 | 0.2059 |
|  | P-tau181 × Time |  | -1.16 | 0.17 | -6.63 | <0.0001* |
|  | Connectome score × P-tau181 |  | -0.07 | 0.37 | -0.19 | 0.9458 |
|  | × Time |  |  |  |  |  |
| MoCA | Connectome score | NCI: 76 | -1.19 | 0.46 | -2.61 | 0.0142* |
|  | P-tau181 | CIND: 146 | -2.08 | 0.66 | -3.15 | 0.0043* |
|  | Connectome score × P-tau181 | DEM: 85 | 0.59 | 1.27 | 0.46 | 0.9255 |
|  | Connectome score × Time |  | -0.31 | 0.12 | -2.57 | 0.0388* |
|  | P-tau181 × Time |  | -1.08 | 0.16 | -6.61 | <0.0001* |
|  | Connectome score × P-tau181 |  | -0.11 | 0.35 | -0.33 | 0.9458 |
|  | × Time |  |  |  |  |  |
| Global cognition | Connectome score | NCI: 75 | -0.22 | 0.06 | -3.87 | 0.0012* |
|  | P-tau181 | CIND: 145 | -0.31 | 0.08 | -3.72 | 0.0009* |
|  | Connectome score × P-tau181 | DEM: 82 | 0.04 | 0.16 | 0.22 | 0.9255 |
|  | Connectome score × Time |  | -0.03 | 0.02 | -2.08 | 0.0779 |
|  | P-tau181 × Time |  | -0.15 | 0.02 | -7.02 | <0.0001* |
|  | Connectome score × P-tau181 |  | -0.08 | 0.05 | -1.67 | 0.2891 |
|  | × Time |  |  |  |  |  |
| Executive function | Connectome score | NCI: 75 | -0.21 | 0.07 | -2.77 | 0.0112* |
|  | P-tau181 | CIND: 145 | -0.21 | 0.11 | -1.98 | 0.0815 |
|  | Connectome score × P-tau181 | DEM: 82 | 0.04 | 0.21 | 0.19 | 0.9255 |
|  | Connectome score × Time |  | -0.04 | 0.02 | -1.72 | 0.1480 |
|  | P-tau181 × Time |  | -0.13 | 0.03 | -4.68 | <0.0001* |
|  | Connectome score × P-tau181 |  | -0.06 | 0.06 | -0.94 | 0.6953 |
|  | × Time |  |  |  |  |  |
| Attention | Connectome score | NCI: 75 | -0.15 | 0.08 | -1.81 | 0.0775 |
|  | P-tau181 | CIND: 145 | -0.17 | 0.12 | -1.46 | 0.1854 |
|  | Connectome score × P-tau181 | DEM: 82 | -0.05 | 0.23 | -0.20 | 0.9255 |
|  | Connectome score × Time |  | -0.05 | 0.02 | -2.51 | 0.0388* |
|  | P-tau181 × Time |  | -0.17 | 0.03 | -5.95 | <0.0001* |
|  | Connectome score × P-tau181 × Time |  | -0.11 | 0.06 | -1.78 | 0.2891 |
| Language | Connectome score | NCI: 75 | -0.20 | 0.07 | -2.74 | 0.0112* |
|  | P-tau181 | CIND: 145 | -0.15 | 0.10 | -1.43 | 0.1854 |
|  | Connectome score × P-tau181 | DEM: 82 | 0.15 | 0.20 | 0.74 | 0.9255 |
|  | Connectome score × Time |  | -0.04 | 0.02 | -2.10 | 0.0779 |
|  | P-tau181 × Time |  | -0.16 | 0.03 | -6.50 | <0.0001* |
|  | Connectome score × P-tau181 × Time |  | -0.11 | 0.06 | -1.98 | 0.2891 |
| Verbal memory | Connectome score | NCI: 75 | -0.19 | 0.07 | -2.83 | 0.0112* |
|  | P-tau181 | CIND: 145 | -0.44 | 0.10 | -4.41 | 0.0001* |
|  | Connectome score × P-tau181 | DEM: 82 | 0.19 | 0.20 | 0.97 | 0.9255 |
|  | Connectome score × Time |  | 0.01 | 0.02 | 0.44 | 0.7182 |
|  | P-tau181 × Time |  | -0.08 | 0.02 | -3.59 | 0.0004* |
|  | Connectome score × P-tau181 × Time |  | 0.01 | 0.05 | 0.21 | 0.9458 |
| Visual memory | Connectome score | NCI: 75 | -0.13 | 0.06 | -2.06 | 0.0481* |
|  | P-tau181 | CIND: 145 | -0.40 | 0.09 | -4.39 | 0.0001* |
|  | Connectome score × P-tau181 | DEM: 82 | 0.17 | 0.18 | 0.94 | 0.9255 |
|  | Connectome score × Time |  | -0.02 | 0.02 | -1.37 | 0.2059 |
|  | P-tau181 × Time |  | -0.11 | 0.02 | -5.31 | <0.0001* |
|  | Connectome score × P-tau181 × Time |  | -0.02 | 0.05 | -0.34 | 0.9458 |
| Visuoconstruction | Connectome score | NCI: 75 | -0.15 | 0.07 | -2.13 | 0.0453* |
|  | P-tau181 | CIND: 145 | -0.12 | 0.10 | -1.16 | 0.2677 |
|  | Connectome score × P-tau181 | DEM: 82 | -0.12 | 0.20 | -0.61 | 0.9255 |
|  | Connectome score × Time |  | -0.03 | 0.02 | -1.47 | 0.2059 |
|  | P-tau181 × Time |  | -0.14 | 0.03 | -5.28 | <0.0001* |
|  | Connectome score × P-tau181 × Time |  | -0.11 | 0.06 | -2.02 | 0.2891 |
| Visuomotor | Connectome score | NCI: 75 | -0.26 | 0.07 | -3.77 | 0.0012* |
|  | P-tau181 | CIND: 145 | -0.20 | 0.10 | -2.05 | 0.0815 |
|  | Connectome score × P-tau181 | DEM: 82 | -0.16 | 0.20 | -0.80 | 0.9255 |
|  | Connectome score × Time |  | -0.003 | 0.02 | -0.16 | 0.8754 |
|  | P-tau181 × Time |  | -0.10 | 0.02 | -4.35 | <0.0001* |
|  | Connectome score × P-tau181 × Time |  | 0.01 | 0.05 | 0.17 | 0.9458 |
| NPI total | Connectome score | NCI: 55 | 1.04 | 0.86 | 1.20 | 0.2299 |
|  | P-tau181 | CIND: 116 | 0.50 | 1.21 | 0.42 | 0.6783 |
|  | Connectome score × P-tau181 | DEM: 69 | -1.58 | 2.36 | -0.67 | 0.9255 |
|  | Connectome score × Time |  | 1.20 | 0.43 | 2.80 | 0.0350* |
|  | P-tau181 × Time |  | 0.97 | 0.54 | 1.78 | 0.0769 |
|  | Connectome score × P-tau181 × Time |  | -0.03 | 1.21 | -0.03 | 0.9797 |

\* indicates significance at $p < 0.05$ , after false discovery rate corrections for multiple comparisons across the twelve cognitive/behavioural measures. Abbreviations: Coeff., coefficient; SE, standard error; Adj. $p$ , $p$ -value adjusted for false discovery rate. CDR, Clinical Dementia Rating; MMSE, Mini-Mental State Examination; MoCA, Montreal Cognitive Assessment; NPI, Neuropsychiatric Inventory; NCI, no cognitive impairment; CIND, cognitive impairment no dementia; DEM, dementia.

where Y_ij_ indicates the outcome variable of interest (e.g., grey matter volume, behavioural or cognitive score) for each participant j at timepoint i, γ indicates the estimated fixed effect coefficients, μ indicates the estimated random effect coefficients, and r_ij_ indicates the residual for each participant j at timepoint i.

P-values for all linear mixed-effects models were corrected for multiple comparisons using false discovery rate (FDR). Linear mixed-effects models were performed using the lmerTest package [52] on R. Visualization of results were performed using either the ggplot2 package [53] on R (for graphs) or the BrainNet Viewer toolbox [54] on MATLAB (for brain maps).

## 3 RESULTS

### 3.1 High cerebrovascular disease burden across multiple MRI markers linked to widespread functional connectivity disruptions

From the PLSC analyses, we identified one significant latent variable (*p* < 0.05) that explained 76.6% of the covariance between CeVD markers and FC (Figure 2A). This latent variable featured positive weights for all CeVD markers, indicating high CeVD burden across all markers. This pattern of high CeVD burden was correspondingly linked to predominantly lower within-network and higher between-network FC in cortical regions, suggesting lower CeVD-related functional segregation of cortical networks. Additionally, the CeVD pattern was associated with lower within-network subcortical connectivity, as well as lower subcortical connectivity to limbic, default, salience/ventral attention and control networks but higher subcortical connectivity to dorsal attention, somatomotor and visual networks. This finding indicated differential effects of CeVD pathology on subcortical connectivity to associative and sensorimotor networks respectively.

**Figure 2:**
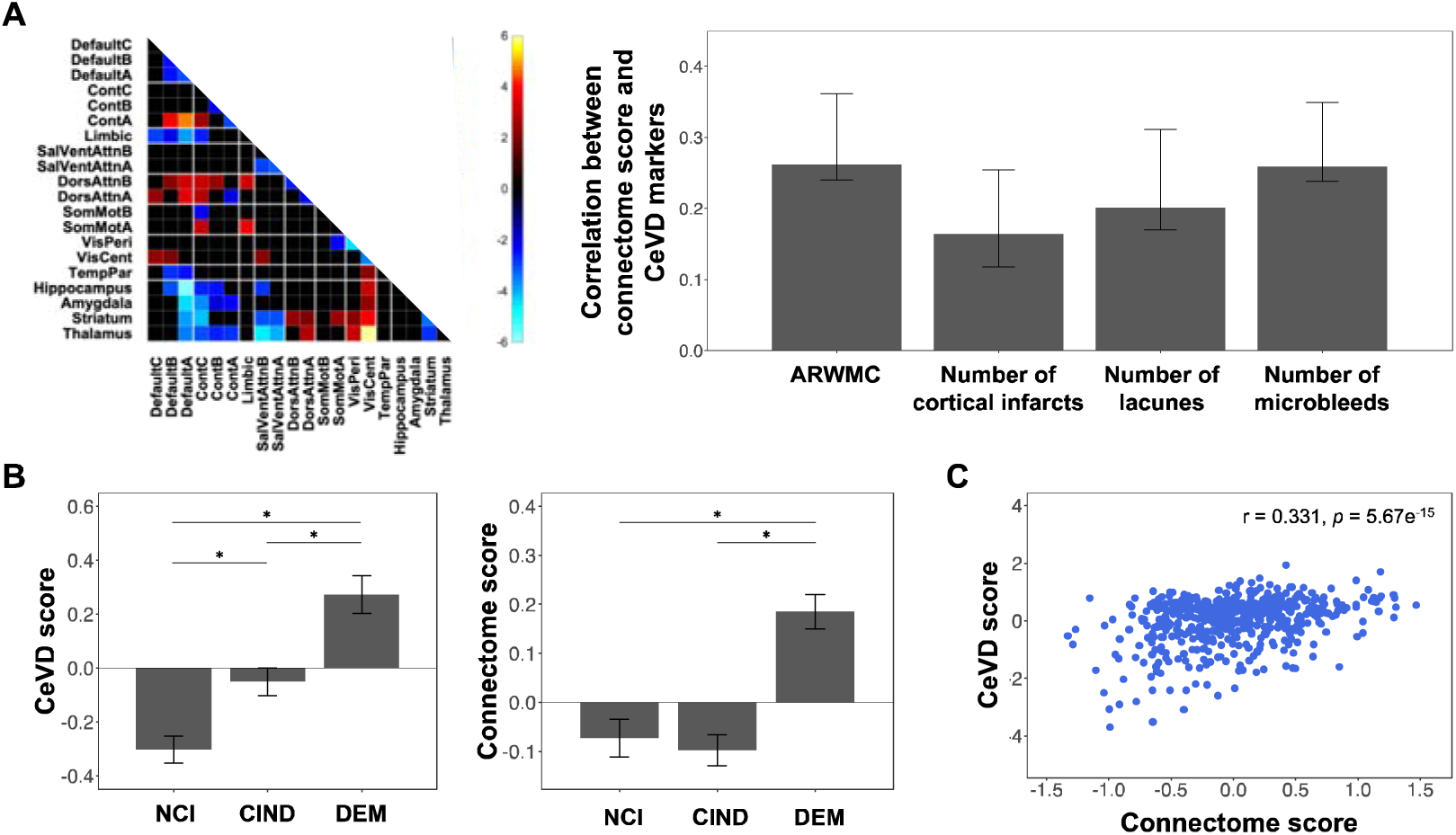
Partial least squares correlation reveals a functional connectome phenotype that is linked to high burden across multiple CeVD markers and characterized by widespread connectivity changes. We identified one significant latent variable that explained 76.6% of the covariance between CeVD markers and FC. (A) Matrix (left) displays the bootstrap ratios (z-scores) of functional connections corresponding to this latent variable. Only significant connections (ratio > 2) are displayed, indicating the stable and reliable connections that contribute to the covariance between CeVD markers and FC. Bar chart (right) denotes the mean correlation between connectome scores of this latent variable and each of the CeVD markers, with error bars indicating 95% bootstrapped confidence intervals. The latent variable featured high CeVD burden across all markers, which was correspondingly associated with predominantly lower within-network but higher between-network FC in cortical regions, as well as lower subcortical connectivity to associative, transmodal networks but higher subcortical connectivity to sensorimotor, unimodal networks. (B) Barplots display mean (± 1 standard error) CeVD and connectome scores for each diagnostic group. * indicate significant pairwise comparisons (*p* < 0.05). Significant differences in CeVD and connectome scores were observed across different diagnostic groups. (C) Scatterplot displays the association between CeVD scores and connectome scores. Abbreviations: FC, functional connectivity; CeVD, cerebrovascular disease; Cont, control network; SalVentAttn, salience/ventral attention network; DorsAttn, dorsal attention network; SomMot, somatomotor network; VisPeri, peripheral visual network; VisCent, central visual network; TempPar, temporal parietal network; ARWMC, age-related white matter changes; NCI, no cognitive impairment; CIND, cognitive impairment no dementia; DEM, dementia.

We further examined how the expression of the CeVD and FC patterns in the latent variable differed across different diagnostic groups by extracting individual CeVD scores and connectome scores pertaining to the latent variable. One-way analysis of variance revealed significant effects of diagnosis on connectome scores (*F*(2, 526) = 20.64, *p* < 0.001), with post-hoc Tukey’s honestly significant difference (HSD) tests showing significantly higher connectome scores in dementia patients compared to CIND (*p* < 0.001) or NCI (*p* < 0.001) patients. Significant effects of diagnosis were similarly observed for CeVD scores (*F*(2, 526) = 19.37, *p* < 0.001), with highest scores in dementia patients and lowest scores in NCI patients (all group differences were significant, *p* < 0.05) (Figure 2B).

### 3.2 Validation of the cerebrovascular disease-related functional connectome phenotype

Similar findings were obtained when PLSC analyses were instead performed on 5 CeVD markers (including number of cortical microinfarcts) and FC in a subset of 436 participants at baseline (Supplementary Figure 2A). Similar findings were also observed when PLSC analyses were performed in a separate dataset comprising a subset of participants (n = 316) with year 2 (two years from baseline) FC and CeVD data (4 markers) (Supplementary Figure 2B). Our findings thus support the reproducibility of the CeVD-related FC phenotype.

### 3.3 Additive effects of the cerebrovascular disease-related functional connectome phenotype and plasma p-tau181 on grey matter volumes at baseline and over time

Having identified a FC pattern that was linked to high pathology across several CeVD markers, we next investigated how this CeVD-related FC pattern was linked to plasma p-tau181 (an AD marker) and downstream outcomes (grey matter volumes, cognitive and behavioural scores). First, we examined the association between connectome scores and plasma p-tau181. CeVD-related connectome scores were not significantly associated with plasma p-tau181, after controlling for age, sex, ethnicity, years of education, handedness, TIV and diagnosis (Supplementary Figure 3).

We then examined if connectome scores and p-tau181 interacted to influence grey matter volumes at baseline and over time using linear mixed effects models. No significant interaction effects of connectome scores and p-tau181 on grey matter volumes at baseline and over time (i.e., connectome score × p-tau181 and connectome score × p-tau181 × time effects) were found, indicating a lack of synergistic effects between connectome score and p-tau181 on grey matter volumes.

However, we observed significant main, divergent effects of connectome score and p-tau181 on grey matter volumes (Figure 3). Specifically, higher CeVD-related connectome scores were associated with widespread lower grey matter volumes at baseline, including hippocampus, default, control, limbic, salience/ventral attention, dorsal attention, somatomotor and visual networks (FDR-adjusted *p* < 0.05) (Figure 3A). There were no longitudinal effects of CeVD-related connectome scores on grey matter volumes (Figure 3B).

**Figure 3:**
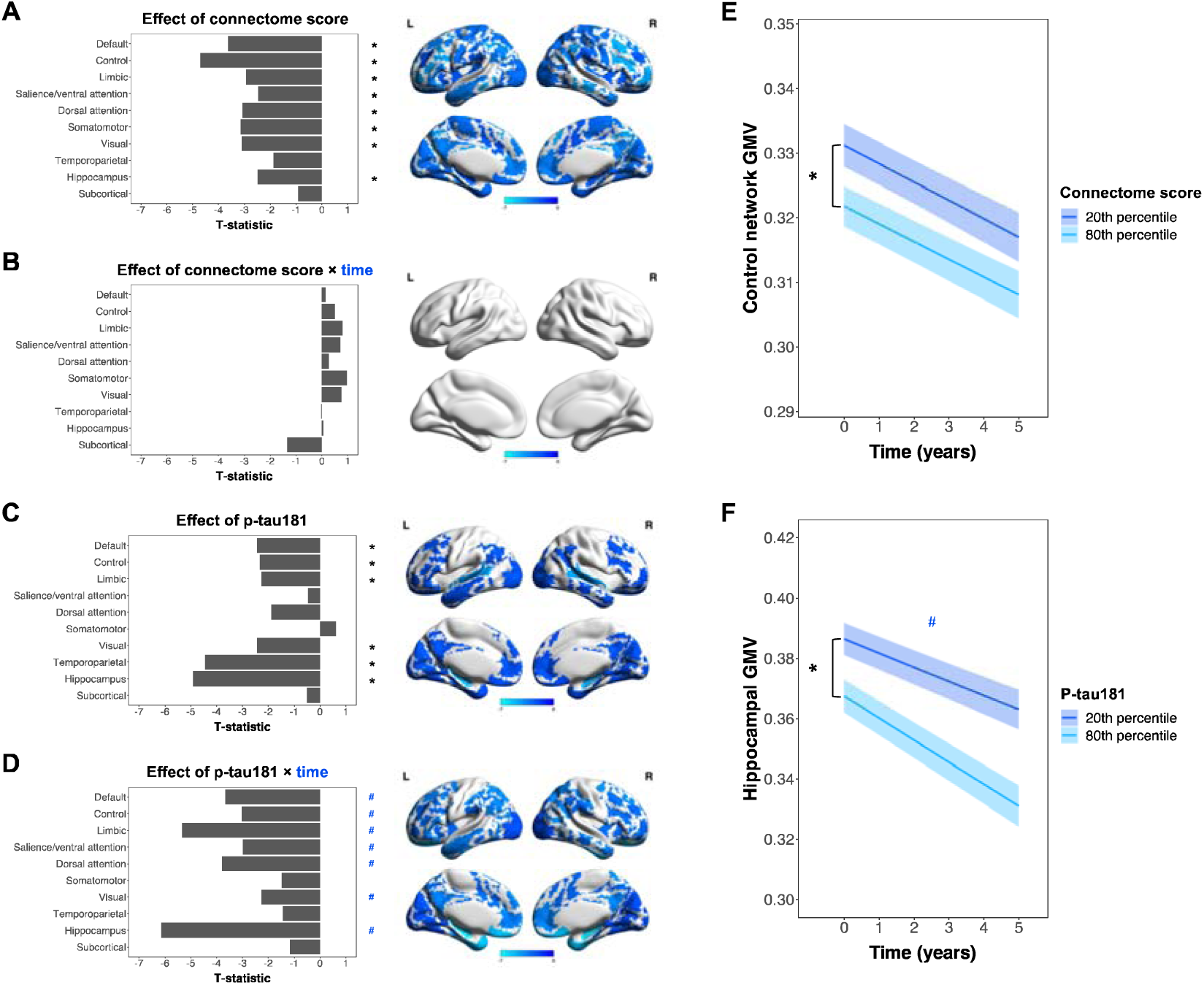
CeVD-related functional connectome phenotype and plasma p-tau181 contribute additively to baseline and longitudinal changes in grey matter volumes. (A-D) Bar plots indicate T-statistic values for the effects of (A) connectome score, (B) connectome score × time, (C) p-tau181 and (D) p-tau181 × time on grey matter volumes across 10 networks. * indicates significant baseline effects (FDR-adjusted *p* < 0.05) while ^#^ indicates significant longitudinal effects (FDR-adjusted *p* < 0.05). Corresponding brain maps display T-statistic values of networks whose grey matter volumes showed significant effects. Lighter blue colours indicate more negative T-statistic values. Additive, but not synergistic effects of connectome score and plasma p-tau181 on changes in grey matter volumes at baseline and over time were observed. Higher connectome scores were associated with widespread lower grey matter volumes across the brain at baseline, but were not associated with grey matter volume changes over time. In contrast, higher p-tau181 was associated with lower baseline grey matter volumes in more localized brain regions, including the hippocampus and default network, but steeper longitudinal declines in grey matter volumes over more widespread areas in the brain. (E-F) The additive effects of connectome score and p-tau181 are further illustrated by representative line plots depicting (E) significant connectome score (FDR-adjusted *p* < 0.05; indicated by *) but non-significant connectome score × time effects on control network grey matter volume, and (F) significant p-tau181 (FDR-adjusted *p* < 0.05; indicated by *) and p-tau181 × time effects (FDR-adjusted *p* < 0.05; indicated by ^#^) on hippocampal grey matter volume. Abbreviations: FC, functional connectivity; CeVD, cerebrovascular disease; FDR, false discovery rate; GMV, grey matter volume.

In contrast, higher p-tau181 was linked to lower grey matter volumes in more localized AD-vulnerable brain regions at baseline, including the hippocampus, default, control, limbic, visual and temporoparietal networks (FDR-adjusted *p* < 0.05) (Figure 3C). Importantly, over time, the effects of p-tau181 on grey matter volumes became more widespread, with higher p-tau181 associated with steeper grey matter volume declines in hippocampus, default, control, limbic, salience/ventral attention, dorsal attention, and visual networks (FDR-adjusted *p* < 0.05) (Figure 3D).

Taken together, our results indicate that p-tau181 and expression of the CeVD-related FC pattern have additive and divergent effects on grey matter volumes both at baseline and over time.

### 3.4 Additive effects of the cerebrovascular disease-related functional connectome phenotype and plasma p-tau181 on cognitive/behavioural performance at baseline and over time

Similar additive and divergent effects of CeVD-related connectome scores and p-tau181 on cognitive and behavioural performance were also observed (Figure 4). We found significant main effects, but not interaction effects, of connectome score and p-tau181 on cognitive and behavioural scores at baseline and over time.

**Figure 4:**
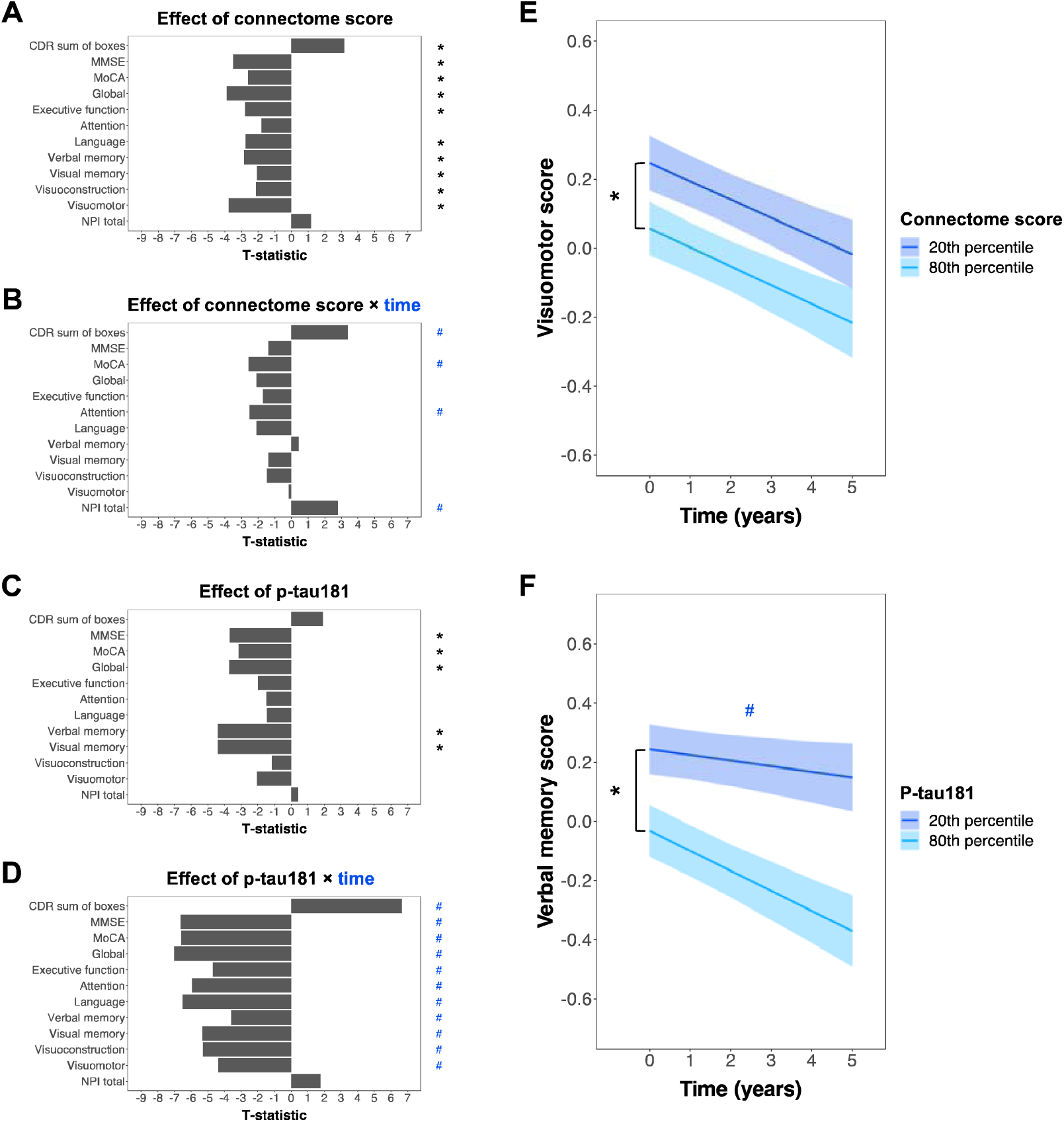
CeVD-related functional connectome phenotype and plasma p-tau181 contribute additively to baseline and longitudinal changes in cognitive and behavioural performance. (A-D) Bar plots indicate T-statistic values for the effects of (A) connectome score, (B) connectome score × time, (C) p-tau181 and (D) p-tau181 × time on cognitive and behavioural scores. * indicates significant baseline effects (FDR-adjusted *p* < 0.05) while ^#^ indicates significant longitudinal effects (FDR-adjusted *p* < 0.05). Additive, but not synergistic effects of connectome score and plasma p-tau181 on baseline and longitudinal changes in cognitive and behavioural performance were observed. Higher connectome scores were associated with widespread lower baseline performance across most cognitive tests, as well as steeper longitudinal declines in MoCA and attention scores and steeper increases in CDR sum of boxes and total NPI scores. By comparison, higher p-tau181 was associated with lower baseline scores in global cognition (MMSE, MoCA and global) and memory (visual and verbal memory) tests, but widespread steeper longitudinal declines across all cognitive tests. (E-F) The additive effects of connectome score and p-tau181 are further illustrated by representative line plots depicting (E) significant connectome score (FDR-adjusted *p* < 0.05; indicated by *) but non-significant connectome score × time effects on visuomotor score, and (F) significant p-tau181 (FDR-adjusted *p* < 0.05; indicated by *) and p-tau181 × time effects (FDR-adjusted *p* < 0.05; indicated by ^#^) on verbal memory score. Abbreviations: FC, functional connectivity; CeVD, cerebrovascular disease; FDR, false discovery rate; CDR, Clinical Dementia Rating; MMSE, Mini-Mental State Examination; MoCA, Montreal Cognitive Assessment; NPI, Neuropsychiatric Inventory.

At baseline, higher CeVD-related connectome scores were linked to lower scores in tests of global functional deterioration (CDR sum of boxes) and global cognition (MMSE, MoCA and global cognition scores), as well as lower scores in six out of seven cognitive domains (executive function, language, verbal memory, visual memory, visuoconstruction and visuomotor performance) (FDR-adjusted *p* < 0.05) (Figure 4A). Over time, higher CeVD-related connectome scores were however associated only with steeper declines in MoCA and attention scores as well as steeper increases in CDR sum of boxes and total NPI scores (FDR-adjusted *p* < 0.05) (Figure 4B).

By comparison, higher p-tau181 was linked to lower performance in global cognition (MMSE, MoCA and global cognition scores) and memory (visual and verbal memory scores) at baseline (FDR-adjusted *p* < 0.05) (Figure 4C). Importantly, higher plasma p-tau181 at baseline was related to widespread steeper declines across all cognitive tests (both global and domain-specific) over time (FDR-adjusted *p* < 0.05) (Figure 4D).

## 4 DISCUSSION

Our study identified an FC phenotype linked to high CeVD burden across four markers in a cohort of participants spanning the AD and CeVD spectrum. This CeVD-related functional connectome phenotype was characterized by widespread FC disruptions, including lower cortical network segregation and opposing subcortico-cortical FC changes in associative and sensorimotor networks. Similar findings were found when the analyses were repeated in a subset of participants with five CeVD markers or in a separate dataset, supporting the reproducibility of this CeVD-related FC phenotype. Further, we showed that expression of this phenotype and p-tau181 contributed additively, but not synergistically, to baseline and longitudinal grey matter volume and cognitive/behavioural changes. Taken together, our study provides evidence that CeVD has global effects on the brain, and that AD and CeVD have additive but not synergistic effects on downstream outcomes.

### 4.1 Presence of a global cerebrovascular disease-related functional connectome phenotype

To our knowledge, our study is the first to examine the effects of multiple CeVD markers on FC using a multivariate approach. Our analyses revealed the presence of a FC phenotype associated with high burden across various CeVD markers and characterized by widespread FC changes across networks, supporting studies showing widespread, global effects of CeVD on FC [9,10]. Specifically, the phenotype featured predominantly lower within-network but higher between-network cortical FC, indicating lower cortical network segregation with higher CeVD burden. This is consistent with a previous study reporting lower local efficiency and clustering coefficient (both segregation measures) in patients with ischemic leukoaraiosis [55]. Interestingly, age has also been widely related to lower network segregation [56], suggesting that CeVD could be a contributive factor to accelerated brain ageing. Supporting this, one study reported a positive association between white matter hyperintensity volumes and SPARE-BA index (a measure of typical age-related brain atrophy pattern) values [57]. Future work could be done to further explore the role of CeVD in brain ageing.

The CeVD-related FC phenotype was also characterized by widespread subcortical FC changes, featuring lower within-network subcortical FC, and lower subcortical FC to transmodal, associative networks (e.g., default, control networks) but higher subcortical FC to unimodal networks (e.g., visual and somatomotor networks). CeVD-related disruptions in subcortico-subcortical and subcortico-cortical connectivity have been demonstrated and related to worse cognitive performance in several studies [11,58,59]. Intriguingly, our finding of opposing subcortico-cortical connectivity changes in associative and sensorimotor networks have been reported in other disorders, such as Parkinson’s disease [60] and schizophrenia [61], reflecting distinct alterations in functionally segregated basal ganglia-thalamocortical circuits [62]. Such shared subcortico-cortical FC changes may thus possibly account for the occurrence of neuropsychiatric symptoms [4,5] and movement impairments [7] in CeVD, although more evidence is required to support this.

### 4.2 Additive and divergent effects of cerebrovascular disease-related brain phenotype and p-tau181 on grey matter volumes and cognitive/behavioural performance

Additionally, our study demonstrated that expression of the CeVD-related FC phenotype (i.e., connectome scores) and p-tau181 showed additive, but not synergistic effects on both baseline and longitudinal changes in grey matter volumes and cognitive/behavioural performance. Connectome scores were also not associated with p-tau181. These findings corroborate previous studies showing differential effects of AD and CeVD on cognitive function [13,17], cortical thickness [13], and FC [15,16], and suggest that the effects of AD and CeVD are additive rather than synergistic.

One step further, closer examination of the main effects of connectome score and p-tau181 revealed the extent of their divergent influence on neurodegeneration and cognition/behaviour both cross-sectionally and longitudinally. Consistent with the involvement of CeVD on cognitive decline across multiple domains [3], higher connectome scores were associated with lower baseline grey matter volumes in most networks as well as lower baseline performance in almost all cognitive domains, indicating global baseline effects of CeVD-related FC changes on neurodegeneration/cognition. In contrast, higher p-tau181 was associated with lower baseline scores only in visual and verbal memory domains, and lower baseline grey matter volumes in more focal brain areas such as the hippocampus, default, limbic and temporoparietal networks. Memory deficits are a key symptom of AD [21], and grey matter atrophy in hippocampal, limbic and default regions have been widely reported in AD patients [63]. Our results also corroborate with a study showing that higher baseline plasma p-tau181 in cognitively impaired individuals was linked to lower baseline grey matter volumes in the precuneus, anterior cingulate and temporal lobes (including hippocampus), as well as greater longitudinal grey matter atrophy in more widespread regions including the precuneus, frontal and temporal lobes [64]. Importantly, our findings support previous work suggesting distinct cognitive profiles in AD and vascular dementia, where AD patients showed impairments in memory, while vascular dementia patients showed impairments across multiple domains including visuoconstruction, attentional and executive abilities [65,66]. Our findings thus highlight the divergent effects of AD and CeVD on neurodegeneration and cognition, with CeVD showing more global effects and AD exhibiting more focal effects.

We similarly observed divergent effects of p-tau181 and CeVD-related connectome score on longitudinal changes in grey matter volumes and cognitive/behavioural performance, although the longitudinal effects differed from the baseline effects. While widespread effects of connectome scores on performance in multiple cognitive domains and grey matter volumes were observed at baseline, higher connectome scores were only linked to steeper longitudinal performance declines in one cognitive domain (attention) and were not associated with longitudinal changes in grey matter volumes. In contrast, p-tau181 was linked to widespread longitudinal changes in grey matter volumes and cognitive performance, as opposed to more focal changes in memory function and grey matter volume at baseline. The differential influence of p-tau181 and connectome score on the trajectories of downstream outcomes could possibly be related to differences in the nature of AD and CeVD pathologies. AD is a progressive disease characterized by gradual spreading of amyloid-β plaques and neurofibrillary tau tangles throughout the brain in stages in a network-specific manner [67], with global deficits in multiple cognitive domains [68] and more widespread grey matter atrophy [69] as the disease progresses. CeVD, by comparison, comprises several heterogeneous MRI markers with different aetiologies [2] and does not have a consistent progression pattern like AD. Correspondingly, individuals with different CeVD profiles have been reported to show different cognitive trajectories [70]. Further, vascular factors have been suggested to influence cognition and neurodegeneration early in the disease process, likely preceding that of AD pathology [71,72]. The potentially earlier influence of CeVD on cognition and neurodegeneration, along with the heterogeneity of CeVD in terms of its progression and effects on cognition, might hence explain the observed widespread baseline effects but limited longitudinal effects of CeVD-related connectome scores on grey matter volumes and cognitive performance. Notwithstanding, higher connectome scores are linked to steeper longitudinal declines in global functional deterioration (CDR sum of boxes) and global cognition (MoCA), underscoring the relevance of CeVD-related connectome scores in predicting future dementia severity and cognitive decline.

Furthermore, higher connectome scores were linked to steeper longitudinal increases in neuropsychiatric symptoms (measured by total NPI score), although no baseline connectome score associations with total NPI score were found. This finding lends some support to evidence implicating CeVD in the occurrence of neuropsychiatric symptoms [4,5]. The limited associations with total NPI score could be attributed to the nature of the CeVD-related FC phenotype, which is associated with high CeVD burden across all markers and characterized by widespread FC changes. Studies have previously linked different CeVD markers to specific neuropsychiatric symptoms [4,5], suggesting CeVD marker-specific, rather than global, effects on neuropsychiatric symptoms. Correspondingly, specific FC changes have been associated with different neuropsychiatric symptoms in prodromal and clinical AD [73]. Future studies could be done to further clarify the global and marker-specific effects of CeVD on FC and consequently neuropsychiatric symptoms.

### 4.3 Limitations

There are some limitations to our study. Firstly, we only considered the severity of CeVD markers in this study. However, studies have suggested that the anatomical locations of CeVD markers may also play a major role in influencing cognition [74,75]. Secondly, we have focused on identifying a brain connectome phenotype of CeVD using FC. However, CeVD-related disruptions in brain structural connectivity [10] have also been identified and could explain variation in cognitive decline that is not explained by FC. Future studies examining the effects of both severity and location of various CeVD markers on functional and structural connectivity as well as their subsequent effects on downstream outcomes could provide an in-depth understanding of how CeVD gives rise to its clinical symptoms via its effects on functional and structural connectivity.

## 5 CONCLUSION

Using a multivariate approach, we found a FC pattern that was linked to high CeVD burden across multiple markers and characterized by widespread cortical and subcortical FC changes. Further, we demonstrated that expression of this phenotype and plasma p-tau181 had divergent effects on baseline and longitudinal changes in grey matter volumes and cognitive/behavioural performance. Our findings suggest that CeVD has widespread, non-marker specific effects on FC, and support the idea that AD and CeVD have additive but not synergistic effects on downstream outcomes. Together, our findings provide insight on the effects of CeVD on distributed functional networks in older adults.

## List of Abbreviations

CeVD: cerebrovascular disease
FC: functional connectivity
AD: Alzheimer’s disease
BOLD: blood-oxygenation-level-dependent
MRI: magnetic resonance imaging
fMRI: functional magnetic resonance imaging
NCI: no cognitive impairment
CIND: cognitive impairment no dementia
DSM-IV: Diagnostic and Statistical Manual of Mental Disorders, Fourth Edition
NINCDS-ADRDA: National Institute of Neurological and Communicative Disorders and Stroke and the Alzheimer’s Disease and Related Disorders Association
NINDS-AIREN: National Institute of Neurological Disorders and Stroke and Association Internationale pour la Recherche et l’ Enseignement en Neurosciences
ARWMC: age-related white matter changes
MMSE: Mini-Mental State Examination
MoCA: Montreal Cognitive Assessment
CDR: Clinical Dementia Rating
NPI: Neuropsychiatric Inventory
WMS-R: Wechsler Memory Scale-Revised
WAIS-R: Wechsler Adult Intelligence Scale-Revised
EDTA: ethylenediaminetetraacetic acid
FLAIR: fluid attenuated inversion recovery
SWI: susceptibility weighted imaging
MRA: magnetic resonance angiography
MPRAGE: magnetization prepared rapid gradient echo
FSL: FMRIB (Oxford Centre for Functional MRI of the Brain) Software Library
AFNI: Analysis of Functional NeuroImages
MNI152: Montreal Neurological Institute 152
VBM: Voxel-based morphometry
CAT12: computational anatomy toolbox
SPM: Statistical Parametric Mapping
DARTEL: Diffeomorphic Anatomical Registration Through Exponentiated Lie Algebra
STRIVE: Standards for Reporting Vascular changes on Neuroimaging
ROI: region-of-interest
PLSC: partial least squares correlation
TIV: total intracranial volumes
FDR: false discovery rate
HSD: honestly significant difference

## Acknowledgements

This study was supported by the Singapore National Medical Research Council (NMRC/OFLCG19May-0035, NMRC/CIRG/1485/2018, NMRC/CSA-SI/0007/2016, NMRC/MOH-00707-01, NMRC/CG/435 M009/2017-NUH/NUHS, CIRG21nov-0007 and HLCA23Feb-0004), RIE2020 AME Programmatic Fund from A*STAR, Singapore (No. A20G8b0102), Ministry of Education (MOE-T2EP40120-0007 & T2EP2-0223-0025, MOE-T2EP20220-0001), and Yong Loo Lin School of Medicine Research Core Funding, National University of Singapore, Singapore.

