## Supplementary Data for "Additive effects of cerebrovascular disease functional connectome phenotype and plasma p-tau181 on longitudinal neurodegeneration and cognitive outcomes"

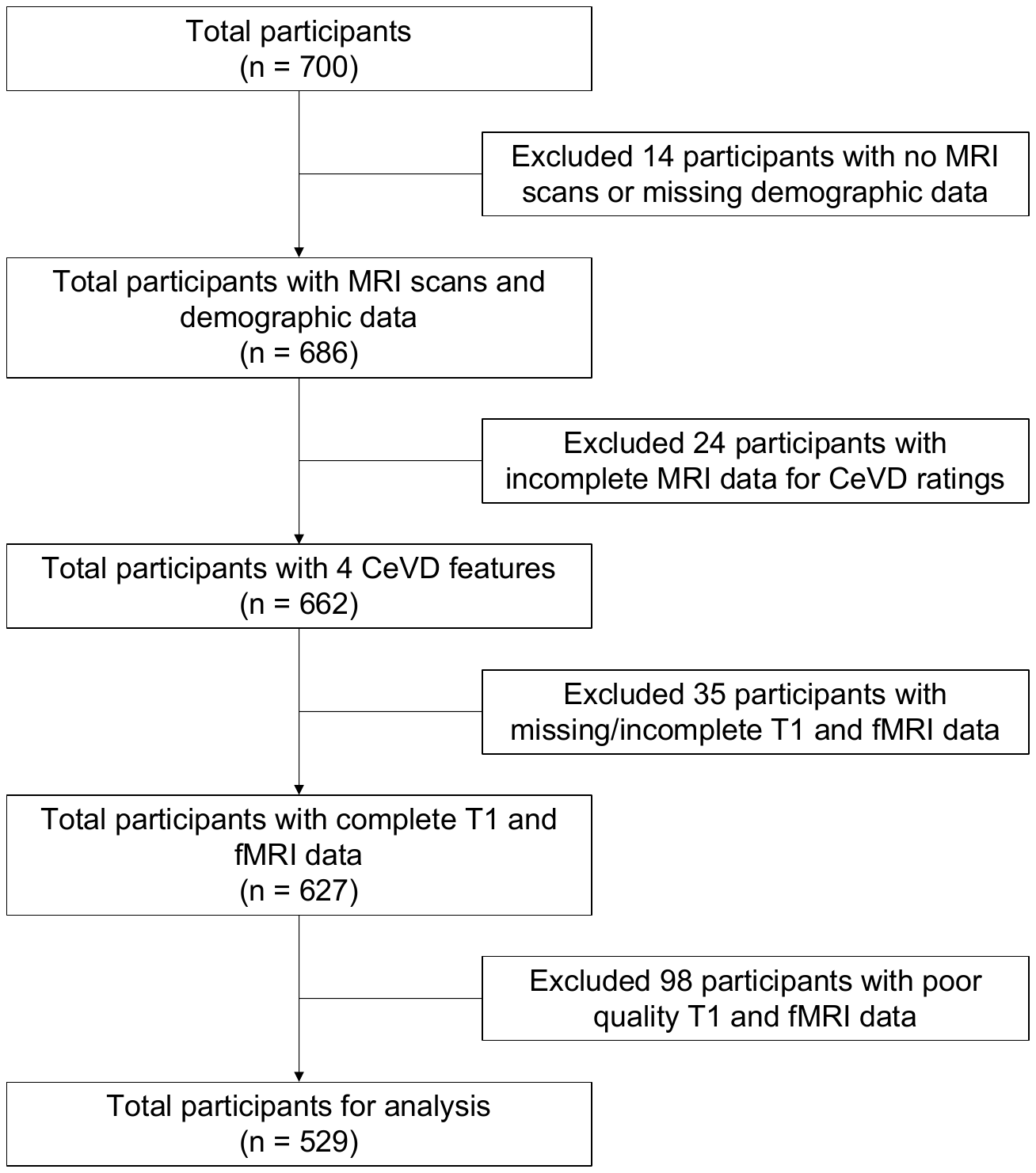

**Supplementary Fig 1: Participant flowchart.** Flowchart illustrates the criteria for selection of participants in the current study. Abbreviations: MRI, magnetic resonance imaging; fMRI, functional magnetic resonance imaging; CeVD, cerebrovascular disease.

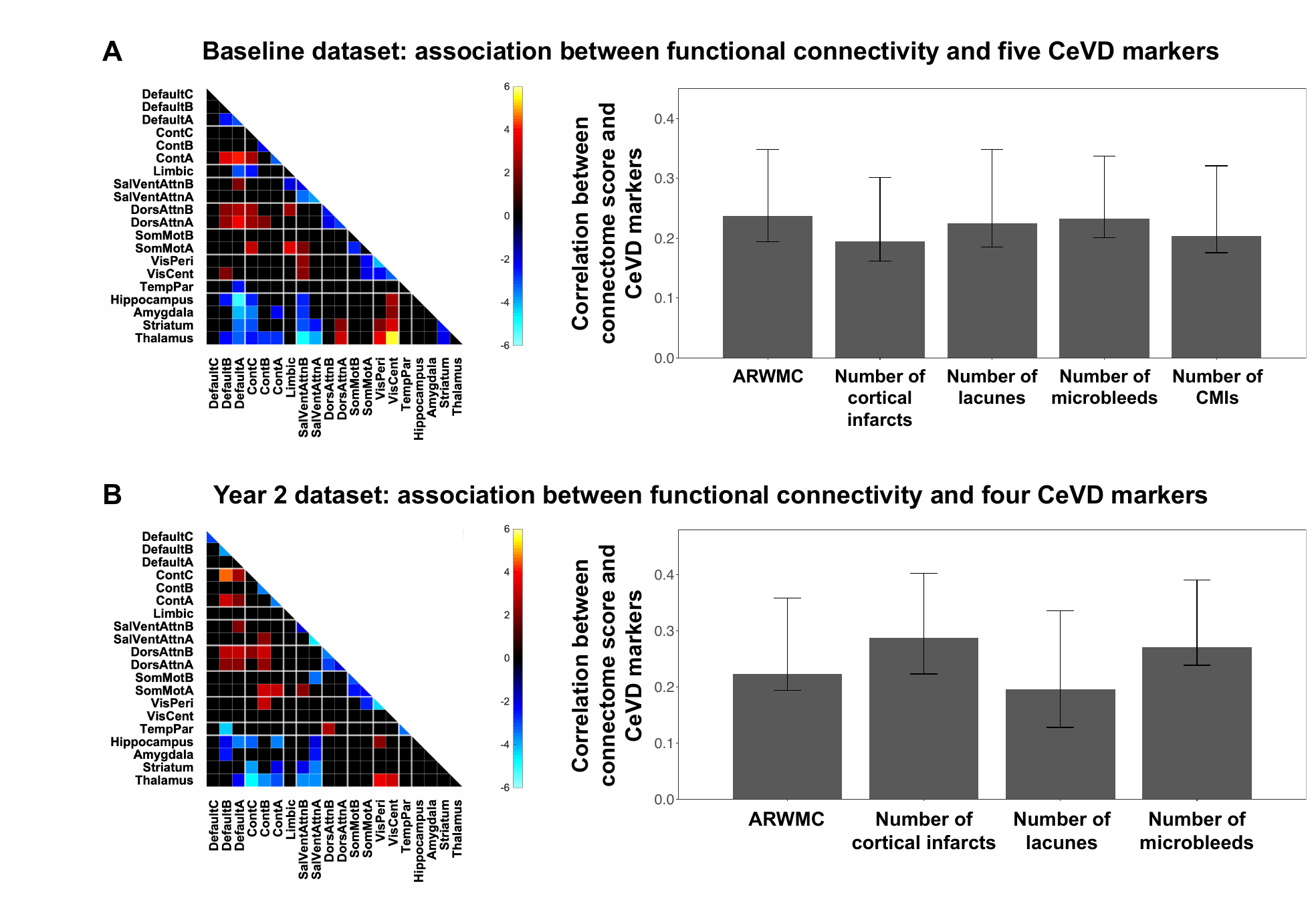
**Supplementary Figure 2: Validation of the CeVD-related functional connectome phenotype.** Partial least squares correlation results for (A) multivariate associations between functional connectivity and five CeVD markers in 436 participants at baseline and (B) multivariate associations between functional connectivity and four CeVD markers in 316 participants at year 2 (i.e., 2 years from baseline). Matrices (left) display bootstrap ratios of significant functional connections (ratio > 2) corresponding to the significant latent variable. Bar charts (right) denote the mean correlation between connectome scores of the significant latent variable and each of the CeVD markers, with error bars indicating 95% bootstrapped confidence intervals. For both sets of analyses, partial least squares correlation identified one significant latent variable that explained (A) 74.4% and (B) 66.3% of the covariance between functional connectivity and CeVD markers respectively. The CeVD and functional connectivity pattern in both sets of analyses mirror that found in the original analyses, indicating the reproducibility of the CeVD-related functional connectome phenotype. Abbreviations: CeVD, cerebrovascular disease; Cont, control network; SalVentAttn, salience/ventral attention network; DorsAttn, dorsal attention network; SomMot, somatomotor network; VisPeri, peripheral visual network; VisCent, central visual network; TempPar, temporal parietal network; ARWMC, age-related white matter changes; CMI, cortical microinfarcts.

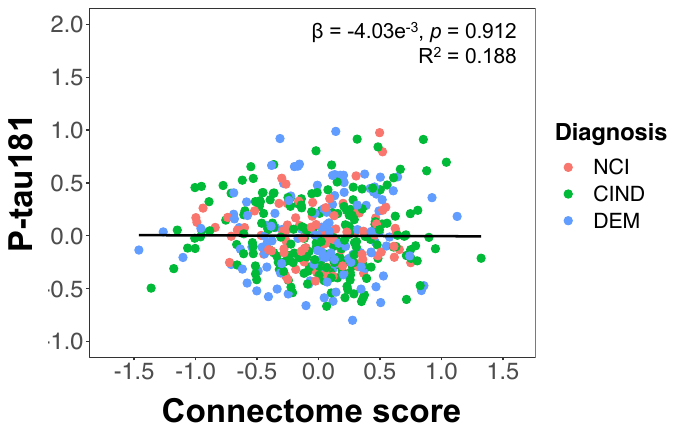

**Supplementary Fig 3: Connectome scores were not associated with plasma p-tau181.** Scatterplot displays the association between plasma p-tau181 and connectome score (n = 413). Connectome scores were not significantly associated with plasma p-tau181, after controlling for age, sex, ethnicity, years of education, handedness, TIV and diagnosis. Abbreviations: CeVD, cerebrovascular disease; TIV, total intracranial volumes; NCI, no cognitive impairment; CIND, cognitive impairment no dementia; DEM, dementia.

**Supplementary Table 1: List of networks**

| **Network** | **Subnetwork** | **Abbreviation** | **Reference** |
| --- | --- | --- | --- |
| Default | Default C | DefaultC | [1] |
|  | Default B | DefaultB |  |
|  | Default A | DefaultA |  |
| Control | Control C | ContC | [1] |
|  | Control B | ContB |  |
|  | Control A | ContA |  |
| Limbic | Limbic | Limbic | [1] |
| Salience/Ventral Attention | Salience/Ventral Attention B | SalVentAttnB | [1] |
|  | Salience/Ventral Attention A | SalVentAttnA |  |
| Dorsal Attention | Dorsal Attention B | DorsAttnB | [1] |
|  | Dorsal Attention A | DorsAttnA |  |
| Somatomotor | Somatomotor B | SomMotB | [1] |
|  | Somatomotor A | SomMotA |  |
| Visual | Peripheral Visual | VisPeri | [1] |
|  | Central Visual | VisCent |  |
| Temporoparietal | Temporoparietal | TempPar | [1] |
| Hippocampus | Hippocampus | Hippocampus | [2] |
| Subcortical | Amygdala | Amygdala | [2] |
|  | Striatum | Striatum | [3] |
|  | Thalamus | Thalamus | Unpublished |

**Supplementary Table 2: Baseline demographic and clinical characteristics for subset of participants with five CeVD markers**

|  | **NCI**  **(n = 93)** | **CIND**  **(n = 198)** | **DEM**  **(n = 145)** | ***p*** |
| --- | --- | --- | --- | --- |
| Age, mean (SD) | 69.8 (7.02)^c,d^ | 72.9 (8.19)^n^ | 74.8 (8.01)^n^ | <0.001 |
| Sex, F/M | 47/46 | 104/94 | 86/59 | 0.3249 |
| Ethnicity, C/M/I/O | 85/2/5/1 | 165/14/17/2 | 116/20/7/2 | 0.0876 |
| Handedness, R/L | 88/5 | 193/5 | 141/4 | 0.4070 |
| Education, mean (SD) | 10.1 (5.2)^c,d^ | 7.5 (4.8)^n,d^ | 5.1 (4.6)^n,c^ | <0.001 |
| MMSE, mean (SD) | 27.3 (1.9)^c,d^ | 24.0 (3.7)^n,d^ | 16.4 (4.6)^n,c^ | <0.001 |
| MoCA, mean (SD) | 25.1 (2.6)^c,d^ | 19.9 (4.7)^n,d^ | 11.6 (4.6)^n,c^ | <0.001 |
| Global CDR, mean (SD) | 0.11 (0.21)^c,d^ | 0.32 (0.24)^n,d^ | 1.28 (0.47)^n,c^ | <0.001 |
| CDR SOB, mean (SD) | 0.14 (0.32)^c,d^ | 0.76 (0.86)^n,d^ | 6.90 (2.83)^n,c^ | <0.001 |
| ARWMC, mean (SD) | 5.35 (3.20)^d^ | 6.35 (3.64)^d^ | 8.68 (4.09)^n,c^ | <0.001 |
| TIV | 1432.0 (146.3)^c,d^ | 1388.4 (147.7)^n^ | 1367.0 (139.8)^n^ | 0.0034 |

^n^ indicates a significant difference from the NCI group, ^c^ indicates a significant difference from the CIND group, ^d^ indicates a significant difference from the DEM group. Group differences in continuous variables were assessed using one-way analysis of variance, while group differences in categorical variables were assessed using chi squared tests. Abbreviations: NCI, no cognitive impairment; CIND, cognitive impairment no dementia; DEM, dementia; F/M, female/male; C/M/I/O, Chinese/Malay/Indian/Others; R/L, right/left; MMSE, Mini-Mental State Examination; MoCA, Montreal Cognitive Assessment; CDR, Clinical Dementia Rating; SOB, Sum of Boxes; ARWMC, age-related white matter changes; TIV, total intracranial volumes.

**Supplementary Table 3: Year 2 participant demographic and clinical characteristics for subset of participants with four CeVD markers**

|  | **NCI**  **(n = 96)** | **CIND**  **(n = 119)** | **DEM**  **(n = 101)** | ***p*** |
| --- | --- | --- | --- | --- |
| Age, mean (SD) | 67.2 (6.14)^c,d^ | 72.0 (8.37)^n,d^ | 77.5 (9.23)^n,c^ | <0.001 |
| Sex, F/M | 56/40 | 57/62 | 66/35 | 0.0312 |
| Ethnicity, C/M/I/O | 85/4/6/1 | 98/12/8/1 | 83/13/3/2 | 0.3411 |
| Handedness, R/L | 91/5 | 114/5 | 97/4 | 0.9028 |
| Education, mean (SD) | 9.7 (5.0)^c,d^ | 8.0 (4.9)^n,d^ | 5.0 (4.7)^n,c^ | <0.001 |
| MMSE, mean (SD) | 27.6 (1.6)^c,d^ | 24.4 (3.3)^n,d^ | 15.6 (4.9)^n,c^ | <0.001 |
| MoCA, mean (SD) | 25.6 (4.2)^c,d^ | 20.6 (4.2)^n,d^ | 11.0 (5.2)^n,c^ | <0.001 |
| Global CDR, mean (SD) | 0.08 (0.18)^c,d^ | 0.25 (0.25)^n,d^ | 1.43 (0.56)^n,c^ | <0.001 |
| CDR SOB, mean (SD) | 0.14 (0.42)^d^ | 0.76 (1.02)^d^ | 8.36 (3.34)^n,c^ | <0.001 |
| ARWMC, mean (SD) | 5.64 (3.39)^c,d^ | 7.27 (3.59)^n,d^ | 9.83 (4.31)^n,c^ | <0.001 |
| TIV | 1391.4 (135.1) | 1412.9 (154.2)^d^ | 1363.4 (129.0)^c^ | 0.0357 |

^n^ indicates a significant difference from the NCI group, ^c^ indicates a significant difference from the CIND group, ^d^ indicates a significant difference from the DEM group. Group differences in continuous variables were assessed using one-way analysis of variance, while group differences in categorical variables were assessed using chi squared tests. Abbreviations: NCI, no cognitive impairment; CIND, cognitive impairment no dementia; DEM, dementia; F/M, female/male; C/M/I/O, Chinese/Malay/Indian/Others; R/L, right/left; MMSE, Mini-Mental State Examination; MoCA, Montreal Cognitive Assessment; CDR, Clinical Dementia Rating; SOB, Sum of Boxes; ARWMC, age-related white matter changes; TIV, total intracranial volumes.
